# The anti-platelet drug cilostazol enhances interrenal steroidogenesis and exerts a scant effect on innate immune responses in zebrafish

**DOI:** 10.1101/2022.12.30.522294

**Authors:** Wei-Chun Chang, Mei-Jen Chen, Yu-Fu Chen, Chih-Wei Chou, Yung-Jen Chuang, Yi-Wen Liu

## Abstract

**Rationale:** Cilostazol, an anti-platelet phosphodiesterase-3 inhibitor used for the treatment of intermittent claudication, is known for its pleiotropic effects on platelets, endothelial cells and smooth muscle cells. However, how cilostazol impacts the endocrine system and the injury-induced inflammatory processes remains unclear.

**Methods:** We used the zebrafish, a simple transparent model that demonstrates rapid development and a strong regenerative ability, to test whether cilostazol influences steroidogenesis, and the temporal and dosage effects of cilostazol on innate immune cells during tissue damage and repair.

**Results:** While dosages of cilostazol from 10 to 100 μM did not induce any noticeable morphological abnormality in the embryonic and larval zebrafish, the heart rate and adrenal/interrenal steroidogenesis in larval zebrafish was increased in a dose-dependent manner. During larval fin amputation and regeneration, cilostazol treatments did not affect the immediate injury-induced recruitment of either neutrophils or macrophages, followed by a subtle yet significant effect on reducing the neutrophil retention at the lesion site.

**Conclusions:** Our results indicate that cilostazol has a significant effect on the growth and endocrine function of steroidogenic tissue; and a limited effect on the neutrophil retention without affecting neutrophil and macrophage recruitment during tissue damage and repair.

## Introduction

Cilostazol is an anticoagulant clinically used to treat peripheral arterial occlusion and thrombotic complications of coronary angioplasty; and for secondary prevention of stroke [1-4]. It is known to increase the intracellular concentration of cyclic adenosine monophosphate (cAMP), a second messenger in the signaling mediated by G protein coupled receptor, by blocking its metabolism through inhibiting the action of phosphodiesterase (PDE) type 3. Both preclinical and clinical studies support the antiplatelet, antithrombosis and vasodilation actions of cilostazol [5, 6]. Owing to these properties, cilostazol has been repurposed as a potential drug for Raynaud’s phenomenon [7]. Moreover, it has been proposed as a potential candidate for treating β-hemoglobinopathies, Alzheimer’s disease and Covid-19 [8-11]. In the dog and human studies, cilostazol treatments lead to typical cardiovascular effects of PDE3 inhibitors such as enhancement in the heart rate, myocardial contractile force and ventricular automaticity [12, 13]. While increasing the concentration of cAMP is known to modulate a multitude of responses in the endocrine and immune systems, how the cilostazol treatment regulates these two systems remain less clear. Understanding how cilostazol impacts the endocrine and immune systems will help us to better address the functions and side effects of cilostazol in the re-purposing studies.

Cilostazol has been found to inhibit secretion of the stress response hormone catecholamine in the bovine adrenal chromaffin cells through inhibiting the Ca2+ movement [14]. However, whether and how cilostazol regulates the function of adrenal cortex remains unclear. PDE2, PDE8 and PDE11 are thought be the major PDEs involved in the cAMP-dependent adrenal steroidogenesis in the mammalian model [15-17]. Nevertheless, a few studies pointed out a possible role of PDE3 and its regulator for modulating the adrenal steroidogenesis. In human adrenocortical NCI-H295 cells, multiple PDE isoforms including PDE3A and PDE3B are down-regulated by p54(nrb)/NONO expression which is essential for adrenocorticotropin response, cAMP production and cortisol synthesis [18]. Moreover, leptin downregulates adrenocorticotropin/cAMP signaling, in NCI-H295 cells, possibly through phosphatidylinositol 3-kinase/Akt and PDE3 [19]. On the other hand, corticosterone synthesis in a neonatal hypoxic-ischemic brain injury rat model is inhibited by granulocyte-colony stimulating factor through JAK2 and PDE3B dependent pathway [20]. As the level of corticosteroid is closely associated with the immune regulation [21], it is of interest to examine whether the PDE3 inhibitor cilostazol affects the function of adrenal cortex.

In the previous *in vitro* preclinical studies, cilostazol has been tested to exert anti-inflammatory effects on effector cells of innate immunity. In the LPS-activated murine RAW264.7 macrophages, cilostazol attenuates cytokine expression, and inhibits high mobility group box 1, NF-kB and PAI-1 through activating AMPK/HO-1 [22, 23]. Cilostazol suppresses LPS-induced PU.1-linked TR4 expression and TLR2-mediated IL-23 production in synovial macrophages isolated from patients with rheumatoid arthritis [24, 25]. In the human monocyte-derived dendritic cells, cilostazol inhibits the production of IL-23 potentially by an AMPK-dependent pathway [26]. However, how cilostazol regulates the *in vivo* behavior of innate immune cells during the process of tissue injury and repair remain unclear.

The tissue transparency, speedy development and a high conserved genome with those in mammals make the zebrafish a robust model for biomedical research. The zebrafish interrenal gland is a functional homolog of mammalian interrenal gland, with highly conserved developmental program and molecular regulation of organ formation [27, 28]. As such, the zebrafish has been established as a teleostean model for hypothalamo-pituitary-adrenal axis and steroidogenesis [29-31]. It is also utilized to examine how glucocorticoids including cortisol, dexamethasone, prednosolone and beclomethasone affect development, immune function and injury-induced regeneration [32-35]. Caudal fin amputation of larval zebrafish has become an *in vivo* model of inflammation and wound repair, due to the unique capability for monitoring the repair process as well as immune cell behavior over a long duration [36, 37]. The sterile wounds are similar to well-controlled surgical wounds which lack complications of infections. It is therefore particularly useful for studying the effects of glucocorticoids on the immune cell recruitment upon tissue injury and repair. Macrophages and neutrophils are both motile phagocytic cells required for the regulation of tissue damage and repair. It is generally accepted that neutrophils dominate the early stages of inflammation, and set the stage for repair of tissue damage by macrophages. Macrophages reside in the tissues while neutrophils typically circulate in the blood, and both can be readily detected and quantified by histochemical or reporter methods in the zebrafish embryo [38]. Zebrafish macrophages and neutrophils start to function during embryonic development, capable of phagocytosis as early as 28 to 30 hours post fertilization (hpf) [39-41].

In this study, we first verified a dose-dependent effect of cilostazol on accelerating the heart rate, further supporting that cilostazol exerts similar pharmaceutical actions in the zebrafish as in mammals. Using the embryonic zebrafish, we then examined whether and how cilostazol affects the development and function of steroidogenic interrenal tissue. Next, we investigated the effect of cilostazol treatments on neutrophils and macrophages during the process of amputation-induced fin regeneration. To our knowledge, this study is the first to demonstrate that cilostazol affects the steroidogenic tissue and endogenous glucocorticoid synthesis, and may therefore impact the glucocorticoid-regulated physiological processes.

## Materials and Methods

### Zebrafish husbandry and egg collection

Zebrafish (*Danio rerio*) were raised according to standard protocols [42]. Embryos were obtained by natural crosses of wild-type and transgenic fish, cultured at 28 °C,and staged as described in [43]. All experimental procedures on zebrafish were approved by the Institutional Animal Care and Use Committee of Tunghai University (IRB Approval NO. 105-28) and carried out in accordance with the approved guidelines.

### Chemical treatments and the heart rate measurement

Stock solutions of 20 mg/mL cilostazol (provided by Otsuka Pharmaceutical) were prepared in DMSO (Sigma). Working solutions were freshly prepared by diluting the stock solutions with aerated egg water. Embryos were placed into these working solutions and cultured at a density of about 30 individuals per 10 ml in a glass beaker. For heart beat measurements, cortisol enzyme-linked immunosorbent assay (ELISA) and whole-mount 3-β-Hydroxysteroid dehydrogenase /Δ5-4 isomerase (3β-Hsd) enzymatic activity assays, the cilostazol treatments commenced immediately after the completion of gastrulation (10 hpf). An assessment of the heart rate is practiced manually by counting heartbeats by a hand-held tally counter. For the tail fin amputation experiments, the cilostazol treatments were started at 1 day post-fertilization (dpf).

### Whole-mount 3β-Hsd enzymatic activity assay

Chromogenic histochemical staining of 3-β-Hsd enzymatic activity was performed on whole embryos essentially according to the described protocol in [44], except that no phenylthiourea treatment for depigmenting was included prior to the assay. Stained embryos from the 3*β*-Hsd activity assay were cleared in 50% glycerol in phosphate-buffered saline (PBS) and subjected to yolk sac removal. The specimens were photographed as ventral flat-mount under Nomarski optics on an Olympus BX51 microscope system.

### Cortisol extraction and ELISA

The treated embryos were harvested at 5 days post fertilization (dpf) for cortisol ELISA assays. The cortisol extraction was performed following the protocol described in [45] with modifications. A group of 30 treated embryos collected as one sample was immobilized by ice-cold water and frozed in liquid nitrogen. The samples were homogenized using a pellet mixer (Dr. Owl) and added with ethyl acetate for the collection of supernatant. The supernatant was evaporized using nitrogen gas, and lipid-containing extracted samples were dissolved in 60 μL of 0.2% BSA in PBS.

For the determination of whole-embryo cortisol levels, a cortisol ELISA kit (Salivary Cortisol Enzyme Immunoassay Kit, Salimetrics) was used according to the manufacturer’s instructions.

Cortisol concentration values were obtained by performing 4 parameter logistic regression of the absorbance readings against a standard curve on a plate reader (TECAN Sunrise).

### Tail fin amputation

Tail fin amputation was performed based on the method described in [46] with modifications. Treated embryos at 3 dpf were anesthetized with 0.025% aminobenzoic acid ethyl ester (tricane, Sigma), and amputated at the posterior edge of the notochord with a 1 mm sapphire blade (World Precision Instruments) on 2% agarose-coated dishes under a Nikon SMZ1500 stereomicroscope. After surgery, the embryos were immediately transferred to freshly prepared solutions for continuous chemical or control treatments.

### Detection and imaging of neutrophils and macrophages

To detect the presence of neutrophils, the staining of myoloperoxidase (mpx) activity by a Peroxidase Leukocyte kit (Sigma) was performed according to the method described in [47] with modifications. Chemical treated and amputated embryos were fixed by 2% paraformaldehyde in PBS (PFAT) for overnight. After washing by PBS containing 0.1% Tween 20 (PBST) and then trizmal buffer containing 0.1% Tween 20 (TT), the staining reaction was carried out in TT buffer containing 0.015% hydrogen peroxide and 1.5 mg/mL peroxidase indicator reagent at 37 °C. The reaction was stopped by washing with PBST and the stained embryos wre post-fixed in 2% PFAT. The fixed embryos were washed with PBST and cleared in 50% glycerol/PBS before being photographed by an Olympus SZX12 microscope.

To detect the macrophage cells recruiting to the amputation site, embryos of *Tg(mpeg1:mCherry)* (from Taiwan Zebrafish Core Facility) were subject to the chemical treatments and amputations, and fixed with 1% PFAT at the indicated time points for temporal analysis. The fixed embryos were washed with PBST and cleared in 50% glycerol/PBS, before being photographed with a LSM510 confocal microscope equipped with LSM 3.5 software (Zeiss). The mCherry fluorescent image marking macrophages was recorded by using 3D z stack and projections.

### Quantification of neutrophils, macrophages and the caudal fin area following amputation

The immune cells near the injury site were quantified by measuring the number of those neutrophils or macrophages present in the counting region as indicated in Fig. 4E and 6E. The tail fin area of fixed and mounted larval samples was measured from the caudal end of the notochord to the edge of the regenerating fin by using Image J software, and expressed as arbituary units.

### Statistical analysis

All statistics were performed with the R studio software (Version 1.4.1106), and graphs were made with GraphPad Prism 8. All quantitative data are presented as mean ± standard error of the mean (SEM). For comparing multiple groups of data, the normality and homogeneity of variances were analyzed by the Shapiro-Wilk test and the Levene’s test. For normally distributed data with homogeneity of variance, ANOVA was used to compare the means of the groups; and the Duncan or Scheffe test for post-hoc analysis. For non-normally distributed data with homogeneity of variance, the Kruskal-Wallis test was used to compare the means of the groups; and the Dunn test for post-hoc analysis. For data that lack homogeneity of variance, the Welch’s ANOVA was used to compare the means of the groups; and the Games-howell test for post-hoc analysis.

## Results

### The cilostazol treatment enhances heart rate in the zebrafish in a dose-dependent manner

In order to test the pharmaceutical actions of cilostazol on the general morphology and development of zebrafish, the embryonic zebrafish were treated with various concentrations of cilostazol from 10 hpf onwards, when the process of organogenesis initiates. Dosages of cilostazol treatments at 10, 30, 50 and 100 μM did not cause noticeable morphological effect on the developing zebrafish, implicating a low toxicity of cilostazol on the growing tissues. Moreover, the heart rate of developing zebrafish was not affected by various concentrations of cilostazol at 29 hours post fertilization (hpf) and 2 days post fertilization (dpf) (Fig. 1). Nevertheless, the cilostazol treatments led to significant increase of heart rate at 3 dpf, a stage when the zebrafish starts a transition from embryos to larvae. At 3 dpf, the heart rate of cilostazol-treated zebrafish was enhanced in a dose-dependent manner from 30 to 100 μM, while the heart rates of 10 and 30μM treatment groups did not show significant difference. This result is consistent with the tachycardia phenomenon seen in cilostazol-treated human adults and the dose-dependent increase of heart rate in healthy dogs treated with cilostazol [48, 49], implicating that the pharmaceutical actions of cilostazol on the developing zebrafish may be similar to those on the mammalian adults.

**Figure 1.**
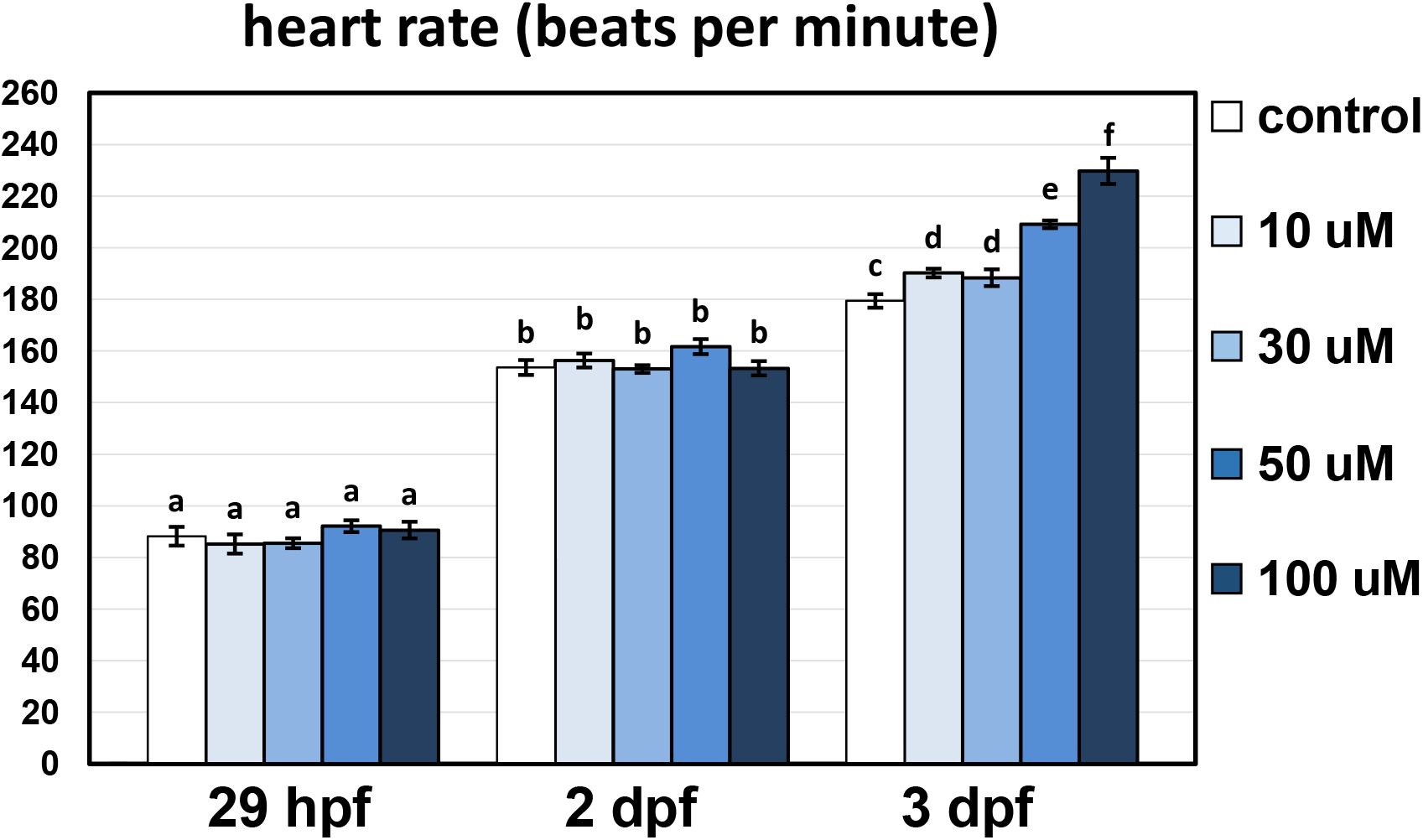
The effect of cilostazol on the developing zebrafish at various stages. The wild-type embryos of zebrafish were treated with increasing concentrations of cilostazol as well as the solvent control from 10 hpf onwards. For each treatment group, 10 embryos were subject to the measurement of heart rate under the microscope at 29 hpf, 2 dpf and 3 dpf respectively. Histograms with different letters above them are significantly different (ANOVA and Duncan’s multiple test, *P* < 0.05). The representative video clips for the heartbeat of 3 dpf embryos in the vehicle control or 100 μM cilostazol-treated groups are shown in Video S1 and S2 respectively.

### The cilostazol treatment leads to an increase of steroidogenic cells as well as cortisol levels in zebrafish

It is known that the activation of protein kinase A (PKA) by cAMP increases the gene expression of StAR (steroidogenic acute regulatory protein), which is crucial for steroidogenesis in the adrenal cells. However, whether and how cilostazol, which enhances cAMP production, affects the cellular physiology of the adrenal gland *in vivo* remains unclear. We therefore used the zebrafish embryo to test whether cilostazol affects development and function of the cortisol-producing steroidogenic tissue. Histochemical staining of 3β-Hsd activity has been routinely used to detect differentiated steroidogenic cells in isolated tissues from mammals as well as teleosts [50, 51], and can be applied to whole-mount embryos and larva of zebrafish [44]. Embryos subject to immersions in cilostazol- or vehicle-containing egg water were assayed at 3 dpf, a stage when the organogenesis of interrenal gland is completed and ready for stress-initiated responses [29, 30]. As pigment formation at the dorsal side of zebrafish pronephros is evident at such stage [52] and often obscures the results of 3β-Hsd staining, we used two low pigment-producing strains, *golden* and *citrine*; and the embryos stained with whole-mount 3β-Hsd activity assay were deyolked and visualized from the ventral side. For both *golden* and *citrine* strains, the embryos treated with cilostazol at 50 and 100 μM respectively demonstrated a significant dosage-dependent increase of steroidogenic interrenal cells, as compared to those treated with vehicle control (Fig. 2A-C).

**Figure 2.**
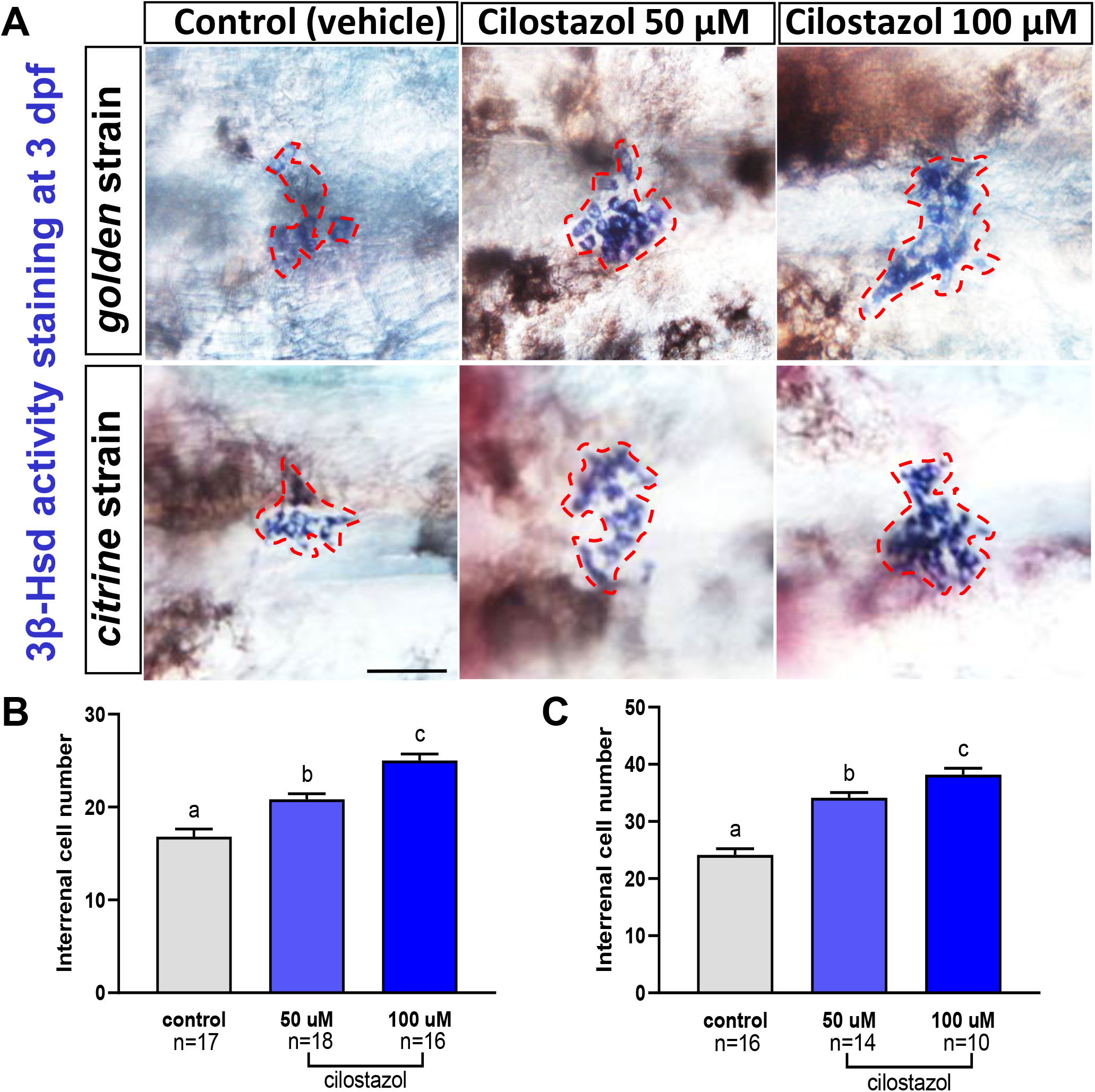
The effect of cilostazol on the morphology and cell number of steroidogenic interrenal tissue in the zebrafish embryo at 3 dpf. The embryos of *golden* and *citrine* strains were treated with cilostazol-supplemented egg water at 50 and 100 μM respectively, or with the vehicle control. The treated embryos were harvested at 3 dpf for whole-mount 3βHsd activity staining. (A) The stained interrenal tissues (highlighted by red dashed lines) were detected on the ventral surface of deyolked embryos with anterior oriented to the left, by using Nomarski microscopy. Scale bar, 50 μm. The number of cells positive for 3βHsd activity staining in the treated *golden* and *citrine* embryos is quantified in (B) and (C) respectively. Histograms with different letters above them are significantly different (ANOVA and Duncan’s multiple test, *P* < 0.05).

Cortisol is the principal steroidogenic hormone for mediating stress response in teleosts [53]. To further estimate whether the increase of differentiated steroidogenic interrenal cells contributes to higher amount of steroidogenic hormone secretion, whole-body cortisol was quantified for larval zebrafish treated with different concentrations of cilostazol or control solutions. As compared to either solvent or blank controls, the cortisol amount in developing zebrafish treated with cilostazol at 10 or 30 μM did not cause increased cortisol levels; yet those treated with cilostazol at 50 or 100 μM led to a significant increase of cortisol amounts at 5 dpf. It is consistent with the increased steroidogenic interrenal cells at 3 dpf old embryos treated with cilostazol at 50 or 100 μM, as compared to those with the solvent control (Fig. 2). While the interrenal steroidogenic cells was increased in a dose-dependent manner at 3 dpf, the cortisol amount at 5 dpf however did not vary between 50 and 100 μM of cilostazol treatments. It is noted that the vehicle control containing a trace amount of DMSO (0.0019%; equivalent to the DMSO amount contained in the 100 μM solution of cilostazol) caused a mild elevation of cortisol level, as compared to the blank control. Nevertheless, cilostazol at 50 and 100 μM led to significantly enhanced steroidogenesis as compared to the solvent control, as evaluated either by counts of interrenal steroidogenic cells (Fig. 2) or by cortisol amounts (Fig. 3).

**Figure 3.**
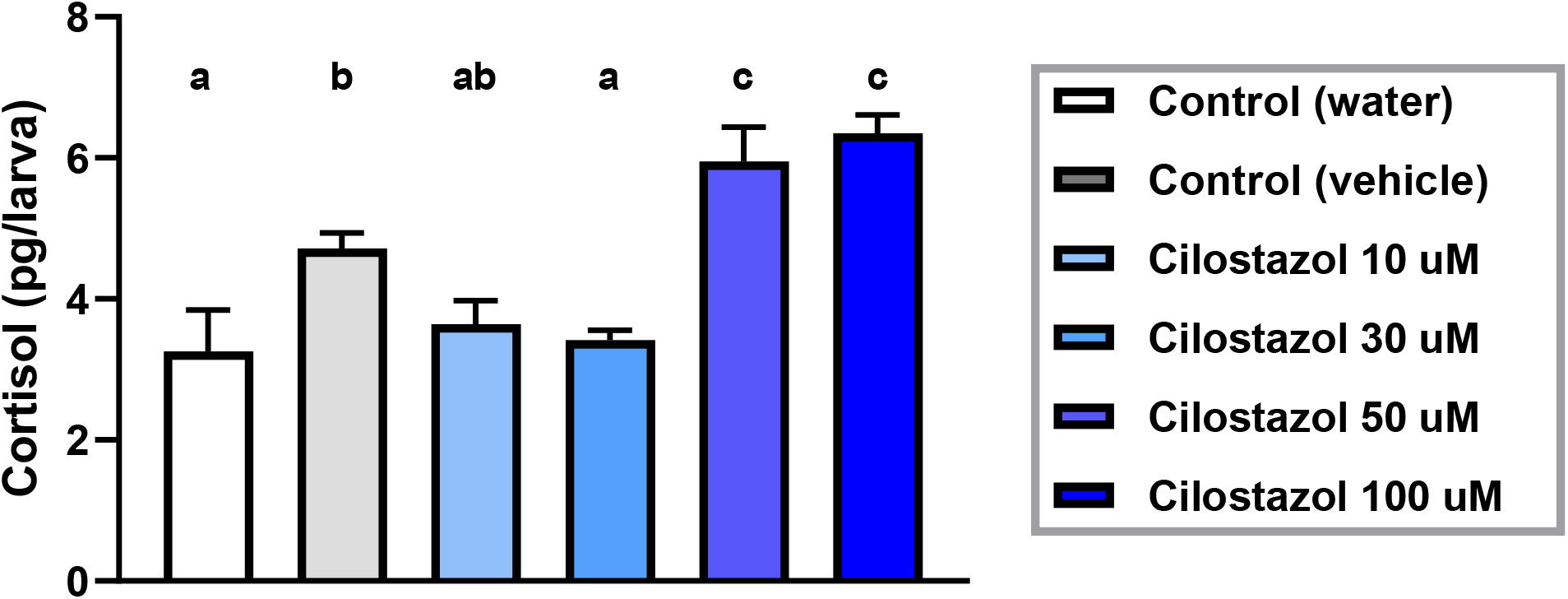
Whole-body cortisol levels after exposure to varying doses of cilostazol as well as controls from 10 hpf to 5 dpf, and measured by ELISA of larval extracts. For each treatment group, 3 samples with each containing 30 larvae were subject to duplicate measurements. Histograms with different letters above them are significantly different (ANOVA and Duncan’s multiple test, P < 0.05).

### The effect of cilostazol treatments on the injury-induced recruitment of neutrophils during zebrafish fin amputation and wound healing

It has been shown in the fin amputation-induced model of inflammation that several synthetic glucocorticoid hormones attenuate the migration of neutrophils toward the wounding site, with that of macrophages unaffected [34, 54, 55]. Whether an elevation of endogenous cortisol levels due to stress or pharmaceutical effects also influence the immune cell accumulation in this wounding-induced inflammation model remains unclear. As our results displayed that cilostazol at 50 and 100 μM enhanced the interrenal steroidogenesis in the zebrafish embryo (Fig. 2 and 3), it is of interest to examine the effect of cilostazol on the behavior of innate immune cells during tail fin amputation and regeneration. The tail amputation was performed on zebrafish embryos at 3 dpf, a stage when neutrophils and macrophages are the two types of leukocytes that constitute the innate immune system. The zebrafish larvae after tail amputation were immediately subject to cilostazol or control treatments (Fig. 4).

**Figure 4.**
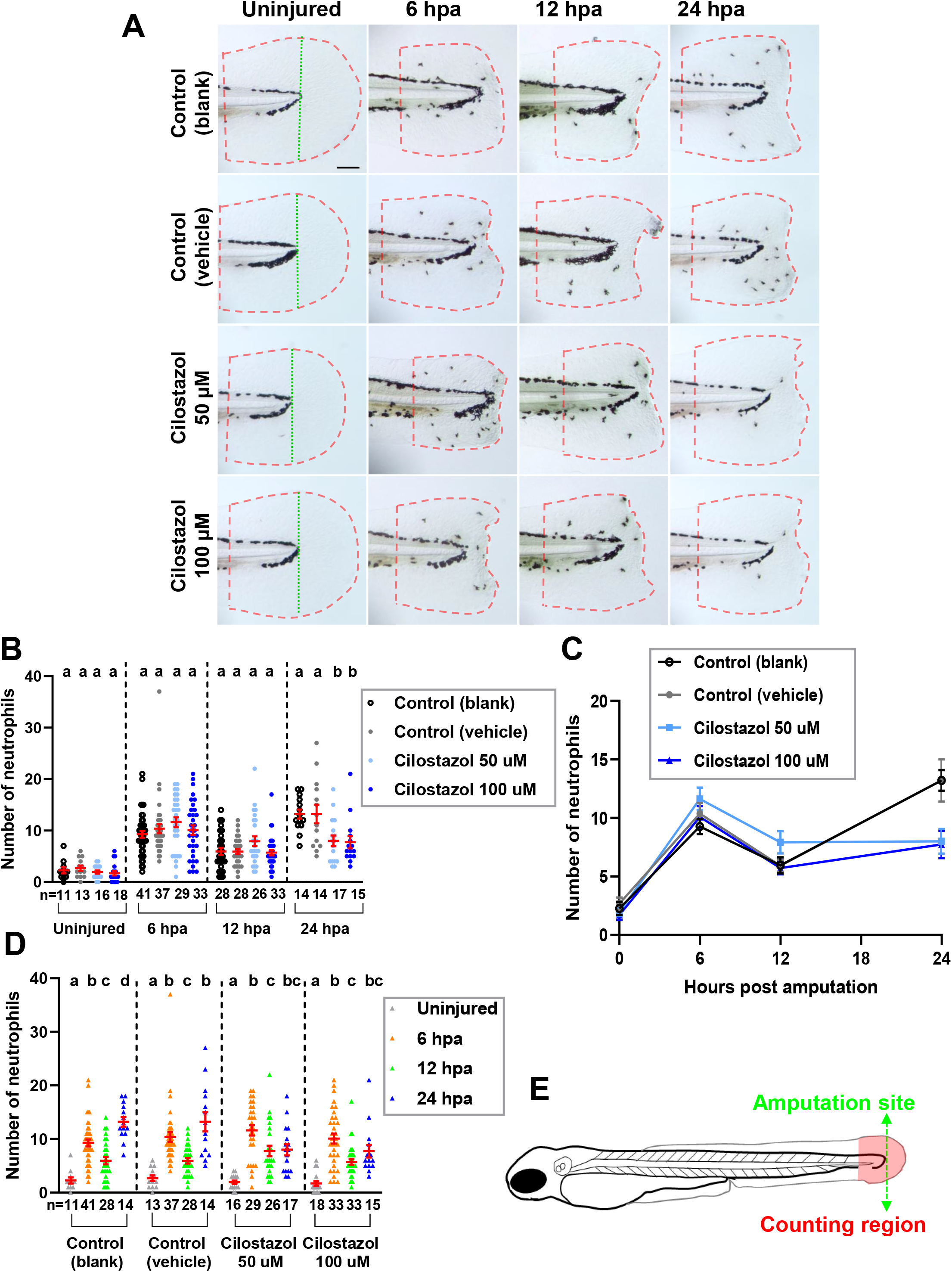
The effect of cilostazol treatment on the accumulation of Mpx-expressing neutrophils during fin amputation and regeneration. (A) The embryonic zebrafish were subject to treatments of cilostazol at 50 and 100 μM respectively, or water and vehicle controls, from 24 hpf onwards to the time of harvest. The amputation of caudal fin fold was performed on the zebrafish embryos at 3 dpf, which were subsequently fixed at 6, 12 and 24 hpa respectively for whole-mount Mpx enzymatic staining. The uninjured fins at 3 dpf were also stained to show the background level of neutrophils prior to their migration to the wounded region. The green dotted line indicates the site of resection, and the area in-between levels of the anterior edge of pigment gap and the posterior edge of regenerating fin, highlighted by red broken lines, was selected for neutrophil quantification. The amputation site and neutrophil counting region are also depicted in the schematic diagram in (E). (B) A comparison of neutrophil accumulation in the selected area of counting as shown in (A), among different treatment groups at the uninjured fins; or at regenerating fins at 6, 12 and 24 hpa respectively. At each time point, differences in the number of neutrophils were compared among various treatment groups. Histograms with different letters above them are significantly different (ANOVA and Duncan’s multiple test, P < 0.05). (C) A line chart showing the temporally dynamic changes of neutrophil accumulation in the wounded fin area, in different treatment groups as shown in (B). (D) A comparison of neutrophil accumulation in the wounded fin area among different time points after fin amputation, in each treatment group as shown in (A). Kruskal-Wallis ANOVA followed by Dunn’s post hoc test was performed for the analysis of water and vehicle control groups. Welch’s ANOVA followed by Games-howell test was performed for the analysis of 50 and 100 μM cilostazol control groups. Histograms with different letters above them are significantly different (P < 0.05).

The neutrophils in the whole-mount embryo were detected by staining the Mpx activity, and the distribution of Mpx activity-positive neutrophils in the tail fin area was analyzed by microphotography at different time points after amputation. The Mpx-activity of neutrophils promoted the formation of brown-black insoluable reaction products that can be readily differentiated from the melanin pigments under Nomarski microscopy (Fig. 4A). Prior to amputation, the neutrophil count was very low in the tail fin area of either control or cilostazol treated embryos. The neutrophil counts in all treatment groups were evidently increased at 6, 12 and 24 hours post amputation (hpa). The randomly distributed pattern of neutrophils in the wounded fin area was consistent with what has been reported for the tail fin-wounding induced inflammation [34], with no obvious difference between control and cilostazol treatment groups. For each time point, a comparison was made among cilostazol treatments at 50 and 100 μM respectively, and blank as well as vehicle controls (Fig. 4B, C). There was no significant difference of neutrophil counts among all treatment groups except for 24 hpa, when the neutrophil counts detected in the blank control (13.2±0.9) and vehicle control (13.2±1.8) groups were significantly higher than those in 50 μM (8.0±1.1) and 100 μM (7.7±1.2) cilostazol treatment groups.

The dynamic changes of neutrophil counts were also analyzed for each treatment group (Fig. 4C, D). In all treatment groups, an evident rise of neutrophil counts from uninjured state to 6 hpa was followed by a slight down-regulation at 12 hpa, suggesting a recruitment of neutrophils toward the wounded tail area in the early phase of tail injury. While the neutrophil counts were further upregulated at 24 hpa in the blank and vehicle control groups; no significant change was revealed in the 50 and 100 μM cilostazol treatment groups. These results suggest that the cilostazol treatment did not affect the recruitment of neutrophils in the early wound, yet may has an effect on the retention of neutrophils during wound healing of the injured caudal fin-fold.

### The effect of cilostazol treatments on the wound healing of zebrafish caudal fin-folds

Apart from evaluating the effect of cilostazol treatments on the neutrophil cell behavior following tail fin injury, we have also checked whether the growth of regenerating wounds was affected by cilostazol. The wound healing process of amputated zebrafish larval fin-fold involves the formation of wound epithelium starting at 6 hpa, and the blastema formation as well as active cell proliferation evident from 24 hpa onwards [56, 57]. Consistent with earlier studies, epithelial cells surrounding the amputation stump contracted and sealed the wound at 6 hpa in the blank control group, and this process was not affected in either blank control or cilostazol treatment groups (Fig. 4A). As the growth of regenerating caudal fin-fold was measured by quantifying the tail area posterior to the caudal end of the notochord, there was an evident increase from 12 to 24 hpa in all treatment groups. Meanwhile, there was no significant difference of quantified caudal fin-fold area among all treatment groups at 6, 12 and 24 hpa. It is therefore concluded that the wound healing in the zebrafish larval fin model is generally not affected by the cilostazol treatment.

### The effect of cilostazol treatments on the injury-induced recruitment of macrophages during zebrafish fin amputation and wound healing

In order to evaluate the influence of cilostazol treatments on the behavior of macrophages in the inflammatory model of zebrafish fin amputation, the *Tg(mpeg1:mCherry)* line was used to detect the presence of macrophages. In zebrafish, the *mpeg1* gene is expressed as early as 28 hpf by both M1 and M2 types of macrophages [37, 58]. The *Tg(mpeg1:mCherry)* embryos were subject to cilostazol or control treatments immediately after tail amputation at 3 dpf, and macrophages in the caudal fin area at different time points after amputation were examined by confocal microscopy (Fig. 6A). In all treatment groups, the macrophage count was low in the uninjured tail fin and evidently increased at 6 hpa, when the macrophages were recruited to and aggregated near the wounding site. While a comparison was made among all treatment groups for each time point, no significant difference of macrophage counts among treatment groups was revealed at 12 and 48 hpf (Fig. 6B, C). At 6 hpa, the 50 μM cilostazol treatment caused a significantly higher macrophage count (37±3.1) as compared to blank control (21.9±2.9) but not to the vehicle control group (35.1±3.8); and the 100 μM cilostazol treatment led to a macrophage count (25.3±4.3) not significantly different from either blank or vehicle control groups. Similarly at 24 hpa, the 50 μM cilostazol treatment caused a significantly higher macrophage count (26.1±2.9) as compared to blank control (12.9±2.3) but not to the vehicle control group (20.1±2.6); and the macrophage count in the 100 μM cilostazol treated group (21.0±3.1) is not significantly different from those of control groups.

While the migration of macrophages toward the wounding site was most evident at 6 hpa in all treatment groups, a retention of macrophages in the caudal fin-fold area was detected at the following stages up to 48 hpa (Fig. 6A, D). In the blank control group, the macrophage count was not significantly decreased at the stages following 6 hpa; and a similar trend was noted in the 100 μM cilostazol treated group. A slight decrease of macrophage counts at the stages following 6 hpa was detected in the vehicle control and 50 μM cilostazol treated groups. In terms of the migration and retention of macrophages in the fin amputation model, our results therefore do not suggest a clear influence by the cilostazol treatments.

## Discussion

To our knowledge, this is the first study to investigate the impact of cilostazol on growth and endocrine function of the steroidogenic tissue. Meanwhile, our experiments have also helped to elucidate the effects of cilostazol on the injury-induced inflammatory and regenerative processes *in vivo*. The results in this study implicate that cilostazol may exert a similar effect on human fetal and adult adrenal glands. The adrenal gland is known to be an organ undergoing constant regeneration, and the adrenal cortex is continuously renewed by stem cells throughout the adult life [59, 60]. As development and renewal of the adrenal cortex are regulated by common regulatory factors such as Hedgehog, Wnt and ACTH/PKA signaling pathways, it is possible that cilostazol administration may affect the progenitor and stem cell populations of the adrenal cortex, thus altering the adrenal function which is critical for homeostasis in the body.

In our study, the heart rate, interrenal tissue growth as well as cortisol secretion in zebrafish were enhanced in the cilostazol treatment experiments. Cortisol is the most important endogenous glucocorticoid in both humans and teleosts. Natural and synthetic glucocorticoids are known to suppress pro-inflammatory macrophages as well as induce anti-inflammatory monocytes and macrophages, thus impeding the expansion of inflammation [21]. Nevertheless, the cortisol regulation of innate immunity in humans can be both pro-inflammatory and anti-inflammatory [61] Interestingly, the LPS-induced inflammatory response in RAW264.7 cells, associated with activated NF-κB and MAPK pathways, can be attenuated by either cortisol [62] or cilostazol [22], implying that an increase of cortisol due to cilostazol treatments may be involved in the suppression of pathogen-induced inflammation.

*In vitro* experiments in earlier studies indicate that cilostazol suppresses the pro-inflammatory cytokines in the LPS-activated murine RAW264.7 as well as human synovial macrophages. [22-24]. However in our *in vivo* experiments, cilostazol exerted no negative effect on either macrophage recruitment or regeneration induced by the fin amputation (Fig. 5, 6). In the fin amputation model, macrophages recruited toward the wound site are essential for the tissue regeneration [38, 63]. The unaffected macrophage accumulation in the wound area of cilostazol-treated fish was therefore consistent with the normal regeneration of amputated fin. In the scenario of sterile injury, the inflammatory response is known to be essential for the repair process [64]. The initial phase of the inflammatory response is marked by the presence of pro-inflammatory cytokines and the infiltration of immune cells into the wound area. In the subsequent resolution phase, the immune cells switch toward a pro-regenerative phenotype with a decline of cytokine secretion; which is followed by the final phase of inflammation marked by tissue regeneration. Our results implicate that the inflammatory response in the early phase of tissue damage is not affected by cilostazol, albeit an enhancing effect on cortisol secretion. It may also suggest that the effects of cilostazol on innate immune cells during the trauma/tissue damage are distinct from those under exogenous pathogen infections. In zebrafish, the innate immune response to microbial infections can be mounted as early as 28 hpf, by the phagocytic activity of neutrophils and macrophages which display conserved transcriptional signatures resembling those in mammals [39]; while the adaptive immunity becomes mature at around 4-6 weeks post-fertilization [65]. Therefore, our results represent the effects of cilostazol on innate immune cells in the absence of T or B cells, during tissue damage. Cilostazol has been tested beneficial in terms of suppressing encephalitogenic T-cell responses and enhancing regulatory T cell activity [66]. In contrast, our results suggest a limited influence of cilostazol on injury-induced inflammation and regeneration in the immunodeficient scenario.

**Figure 5.**
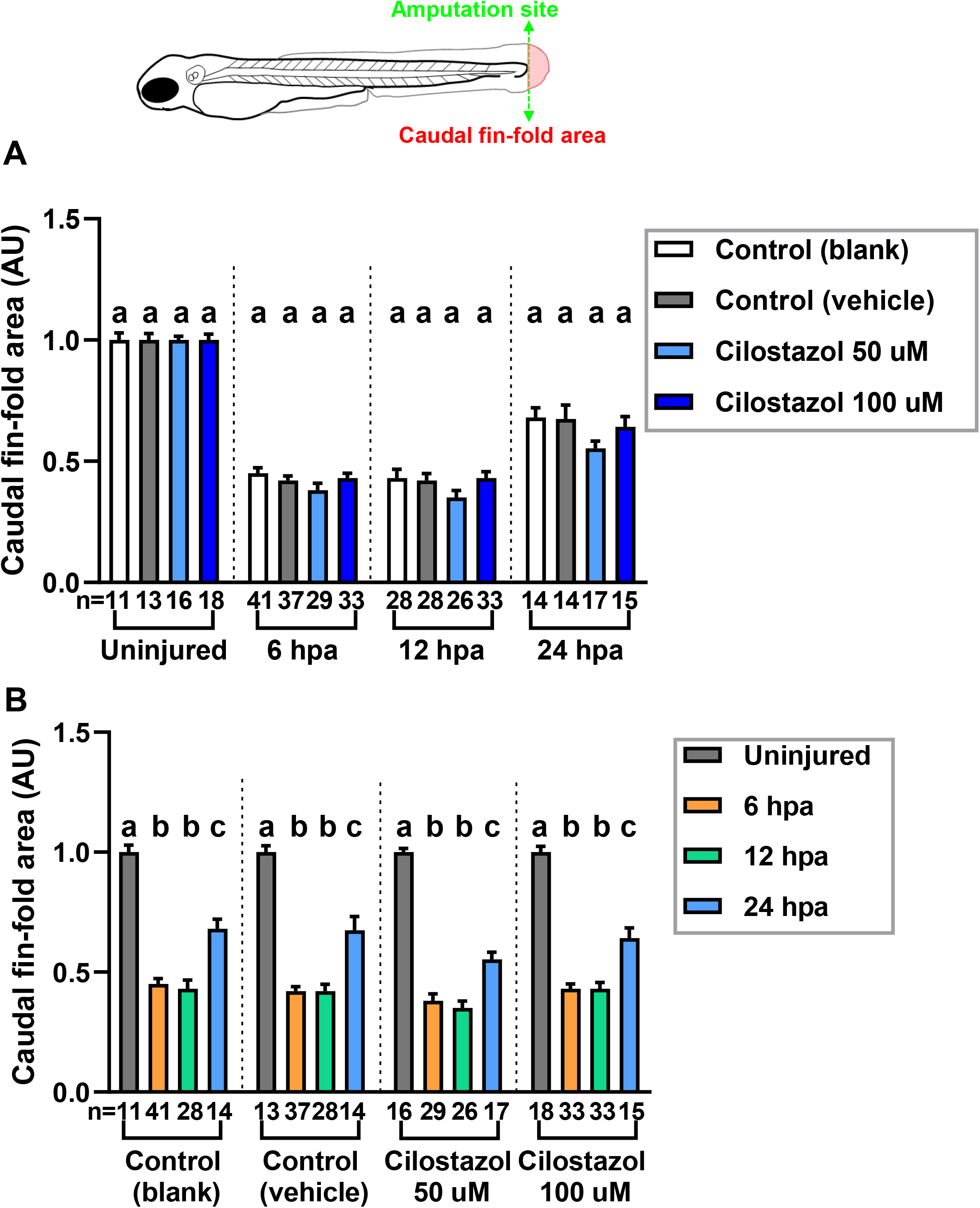
The effect of cilostazol on the regeneration of caudal fin fold zebrafish. The embryos subject to the cilostazol treatments and fin amputation as in Fig. 4 were also analyzed to evaluate whether cilostazol affects the regeneration of fin fold post-injury. The regenerated fin area is defined as the surface area of the fin fold posterior to the amputation site, as viewed laterally. (A) The regenerated caudal fin area of the injured embryos under different treatments are quantified and compared at 6, 12 and 24 hpa respectively, following the amputation at 3 dpf. The uninjured fin areas posterior to the hypothetical site of resection are also quantified and compared among different treatments at 3 dpf. ANOVA analysis followed by Scheffé post hoc test was performed for multiple comparisons among different treatment groups of uninjured fins, and injured fins at 24 hpa. Histograms with different letters above them are significantly different (P < 0.05). Kruskal-Wallis ANOVA followed by Dunn’s post hoc test was performed for comparisons among different treatment groups of injured fins at 6 and 12 hpa. Histograms with different letters above them are significantly different (P < 0.05). (B) The regenerated fin area for each type of cilostazol or control treatments is compared among 6, 12 and 24 hpa following amutation at 3 dpf; and contrasted with the caudal fin area of uninjured embryos at 3 dpf. Kruskal-Wallis ANOVA followed by Dunn’s post hoc test was performed for the analysis of water control group. Welch’s ANOVA followed by Games-howell test was performed for the analysis of vehicle control and 50 μM cilostazol treatment groups. ANOVA analysis followed by Scheffé post hoc test was performed for the analysis of 100 μM cilostazol treatment group. Histograms with different letters above them are significantly different (P < 0.05).

**Figure 6.**
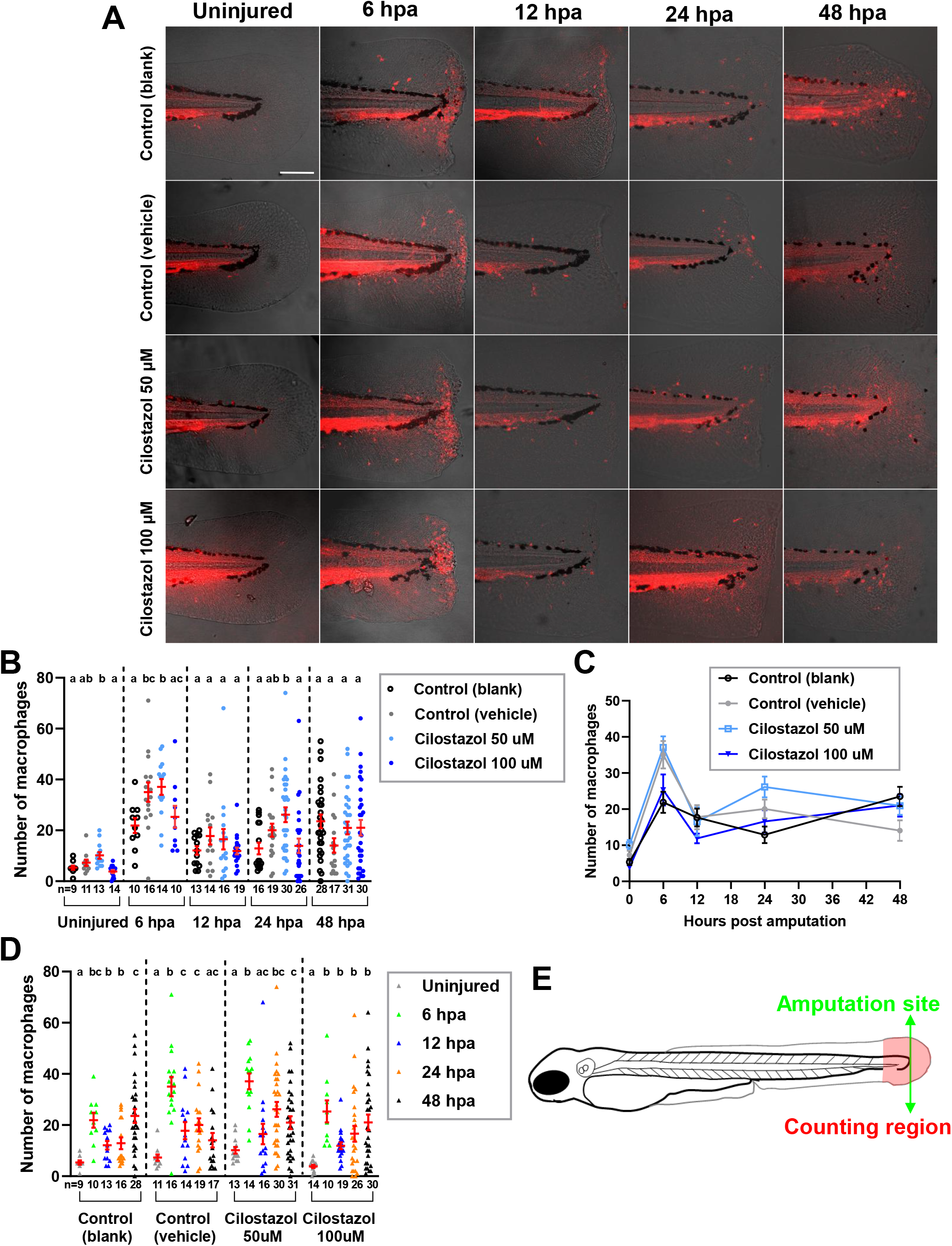
The effect of cilostazol treatment on the accumulation of Mpeg1-expressing macrophages during fin amputation and regeneration. (A) The embryos of *Tg(mpeg1:mcherry)* line were subject to treatments of cilostazol at 50 and 100 μM respectively, or water and vehicle controls, from 24 hpf onwards to the time of harvest. The amputation of caudal fin fold was performed on the embryos at 3 dpf, which were subsequently fixed at 6, 12, 24 and 48 hpa respectively for macrophage quantification. The uninjured fins at 3 dpf in different treatment groups were also collected for comparisons. The green dotted line indicates the site of resection, and the area in-between levels of the anterior edge of pigment gap and the posterior edge of regenerating fin, highlighted by white broken lines, was selected for neutrophil quantification. The amputation site and neutrophil counting region are also depicted in the schematic diagram in (E). (B) A comparison of macrophage accumulation in the selected area of counting as shown in (A), among different treatment groups at the uninjured fins at 3 dpf; or at regenerating fins at 6, 12 and 24 hpa respectively. Kruskal-Wallis ANOVA followed by Dunn’s post hoc test was performed for the multiple comparisons among different treatment groups at 0, 12, 24 and 48 hpa. ANOVA analysis followed by Duncan’s test was performed for multiple comparisons among different treatment groups at 6 hpa. Histograms with different letters above them are significantly different (P < 0.05). (C) A line chart showing the temporally dynamic changes of macrophage accumulation in the wounded fin area, in different treatment groups as shown in (B). (D) A comparison of macrophage accumulation in the wounded fin area among different time points after fin amputation, in each treatment group as shown in (A). Welch’s ANOVA followed by Games-howell test was performed for the analysis of water control as well as 50 and 100 μM cilostazol treatment groups. Kruskal-Wallis ANOVA followed by Dunn’s post hoc test was performed for the analysis of vehicle control group. Histograms with different letters above them are significantly different (P < 0.05).

Although it is generally uncertain whether cilostazol is safe and effective for use in children, a clinical trial testing the effects of cilostazol for the treatment of juvenile Raynaud’s Phenomenon, a blood vessel disorder in children, has been completed with no significant beneficial effects discovered [67]. On the other hand, cilostazol has been tested effective in a rat model of prenatal valproic acid-induced autism spectrum disorder [68] and hence proposed as an adjunctive therapy for the children with autism spectrum disorder [69]. However, our study suggests potential adverse endocrine effects of cilostazol for use in developing young patients, which need to be carefully assessed. The maximal level of cortisol increase caused by the cilostazol treatment was 1.36 fold as compared to the solvent control (Fig. 3). On the other hand, direct early exposure of zebrafish embryos to the cortisol-containing medium, in the study by Hartig et. all, leads to a maximal 1.3 fold elevation of whole-body cortisol amount; which culminates in upregulated basal expressions of pro-inflammatory genes during the later adult stage accompanied by defective adult tailfin regeneration [32]. In human and animal studies, early life stress is known to cause long lasting effects on developing and function of the brain, even causing persistent mental disorders such as anxiety symptoms [70]. Early life stress in the form of childhood adversity is linked to epigenetic regulation of glucocorticoid receptor in the brain, as well as altered neuronal gene expressions due to sustained DNA hypomethylation [71, 72]. In the zebrafish, long lasting anxiety following early life stress is mediated by cortisol and glucocorticoid signaling [73]. The studies above suggest that the hyperactivated glucocorticoid signaling during early life causes persistent effects on the neuronal as well as immune functions lasting to adulthood. Therefore, the results of our study implicate that exposure to cilostazol during early development may lead to pro-inflammatory adults with changes in mental function, which should be considered in the treatment of young patients.

## Acknowledgements

The authors would like to thank Otsuka Pharmaceutical for providing pure cilostazol substance. We would also like to acknowledge the technical services provided by the Taiwan Zebrafish Technology and Resource Center of the National Core Facility Program for Biotechnology, National Science Council, Taiwan. This work was supported by National Science and Technology Council (NSTC) 106-2313-B-029-002-MY3 and 109-2313-B-029-002-MY2 (Taiwan).

## Figure legends

**Video S1**. The heartbeat of a representative 3 dpf embryo in the vehicle control group.

**Video S2**. The heartbeat of a representative 3 dpf embryo in the 100 μM cilostazol-treated groups.

