## Supplemental videos 1, 2 for "The anti-platelet drug cilostazol enhances interrenal steroidogenesis and exerts a scant effect on innate immune responses in zebrafish"

**Video S1.** The heartbeat of a representative 3 dpf embryo in the vehicle control group.

https://reurl.cc/mZQ84G

**Video S2.** The heartbeat of a representative 3 dpf embryo in the 100 μM cilostazol-treated groups.

https://reurl.cc/AylVLp
